# Chikungunya Outbreak in Bangladesh (2017): Clinical and hematological findings

**DOI:** 10.1101/639872

**Authors:** Saeed Anwar, Jarin Taslem Mourosi, Fahim Khan, Mohammad Ohid Ullah, Olivier M. Vanakker, Mohammad Jakir Hosen

**Author notes:** These authors have contributed equally to this work (Saeed Anwar, Jarin Taslem Mourosi, Fahim Khan). Correspondence: Prof. Dr. Mohammad Jakir Hosen, Department of Genetic Engineering and Biotechnology, School of Life Sciences, Shahjalal University of Science and Technology, Sylhet 3114, Bangladesh.

## Abstract

A massive outbreak of Chikungunya occurred in Bangladesh during the period of April-September, 2017 and over two million people were at risk of getting infected by the virus. A prospective cohort of viremic patients was constituted and analyzed to define the clinical, hematological and long-term aspects of this outbreak. A 35-day long comprehensive survey was conducted in two major, neighboring cities, Dhaka and Mymensingh. One-hundred and eighty-seven clinically proven Chikungunya cases were enrolled in the cross-sectional cohort study. Additionally, a smaller group of 48 Chikungunya patients was monitored for post-infection effects for 12 months. Clinical data revealed that a combination of fever and arthralgia (oligoarthralgia and/or polyarthralgia) was the cardinal hallmark (97.9% of cases) of the infection. Hematological analysis showed that, irrespective of age groups, hemoglobin level significantly decreased and erythrocyte sedimentation rate remarkably increased in Chikungunya confirmed patients. However, the majority of the patients had a normal range of whole WBC and platelet counts; RBC counts for mid aged (40 – 60 years) and senior (61+ years) patients (especially in the females) were beyond the reference values. The post-infection study revealed that children had an early recovery from the infection compared to the adults. Moreover, post-infection weakness, successive relapse of arthralgic pain and memory problems were the most significant aftereffects, which had an impact on daily activities of patients. This study represents a comprehensive overview of clinical and epidemiological features of the 2017 outbreak of Chikungunya in Bangladesh as well as its chronic outcomes till the 12^th^ month. It provides insights into the natural history of this disease which may help to improve management of the Chikungunya patients.

**Author summery:** The clinical profile, epidemiology and the economic impacts during the acute phase of Chikungunya infection has been studied quite rigorously. However, studies regarding the hematological features and chronic consequences are very limited. In this study, a dataset of 187 clinically proven chikungunya patients were analyzed for the clinical and hematological features at acute phase of the infection. Additionally, the long-term consequences till month 12 after the infection were studied for a smaller group of 48 patients. Clinical data revealed that a combination of fever and joint pain (arthralgia) was the cardinal hallmark in the acute phase of the infection. Hematological analysis showed that, hemoglobin levels of the patients were significantly reduced and erythrocyte sedimentation rate increased remarkably. Also, RBC counts for mid-aged and older patients were beyond the reference values. The post-infection consequence study unveiled that children recovered better from the infection compared to the adults. Further, post-infection weakness, successive relapse of joint pain and memory problems were the most significant aftereffects. Overall, the infection had moderate to severe impact on daily activities of the respondents. This study provides insights into the clinical and hematological aspects of Chikungunya infection during the acute phase as well as describes an account for its chronic outcomes which puts forward to the knowledge for clinicians and epidemiologists regarding the infection diversity and to help improved patient management.

## Introduction

Chikungunya is a neglected tropical disease, usually endemic to Africa, Southern and Southeast Asia. This disease is caused by the Chikungunya virus (ChikV), a classical arbovirus which possesses a single stranded positive-sense RNA genome that is transmitted to human through the bites of infected female *Aedes* mosquitoes, predominantly by *Aedes aegypti* and *Aedes albopictus* [1-4]. In classical Chikungunya, after a short incubation period of about 1 to 5 days, acute commencement of fever and polyarthralgia, most eminently affecting the limb extremities, are the most frequently reported clinical signs in 72-97% of cases [1, 3, 5]. Other symptoms include skin rash, headaches, back pain, myalgia and nausea [2, 6, 7]. Joint pain can often be severe and may remain indefatigable for weeks to years. The most severe forms of the disease which have been reported are associated with neurological, cardiovascular, hepatic, dermatological or respiratory symptoms along with miscarriages and neonatal infections [8-14]. Despite the fact that only few patients require hospitalization, there have been a few reports of fatalities due to this infection [15-18]. The clinical manifestations of Chikungunya are often confused with Dengue, especially in regions where both diseases can have outbreaks.

Till now, outbreaks of Chikungunya have been reported in more than 60 countries [7]. The first Chikungunya outbreak in Bangladesh was recognized in 2008 in two villages of northwestern Bangladesh [19]. Later, in November 2011, another Chikungunya outbreak was reported in Dhaka [20]. The most dangerous outbreak of Chikungunya in Bangladesh was reported in April – September 2017, when a huge number of positive cases were reported from 23 districts of the country; 13000 clinically confirmed cases were documented in the city of Dhaka alone [21-27]. The ChikV from the 2017 outbreak in Dhaka was found to be genetically distinct from the strain found in the previous outbreak, Bangladesh/0810atw [28]. Phylogenetic analysis revealed that the outbreak strains constituted a new cluster within the Indian Ocean clade, suggesting that they are novel variants [29]. Together with variability in symptoms, 83% of patients in Dhaka also had low to very low overall quality of life, and ∼30% patients had ambulatory problems due to severe arthropathy [30]. However, the impact of ChikV on hematological parameters and its long-term effects have not yet been studied. In this study, hematological parameters were assessed during the Chikungunya 2017 outbreaks in a cohort of 187 patients; a subgroup of these was continuously followed afterwards to better understand the long-term effects.

## Materials and Methods

### Patient recruitment, sample collection and data analysis

During the period of June 30, 2017 to August 4, 2017, 187 clinically confirmed, RT-PCR or serological test positive patients from Dhaka and Mymensingh were recruited randomly in this study (**Fig** 1). A cross-sectional study was done to investigate the clinical, biochemical and hematological profiling, and a long-term follow-up was conducted to understand the after effect of Chikungunya on the quality of life. Only clinically proven Chikungunya patients were included; patients with respiratory and cardiovascular symptoms, previous report of arthralgia, any kind of arthritis, rheumatism, any major recent injuries or blood disorders were excluded from the study. Patients with proven evidence of previous or present infection by Dengue virus were also not included in the study. Among the 187 enrolled patients, 48 were found willing to follow a long-term monitoring scheme of 12 months (Long term consequence assessment group; LCA) (**Fig** 1).

**Fig 1:**
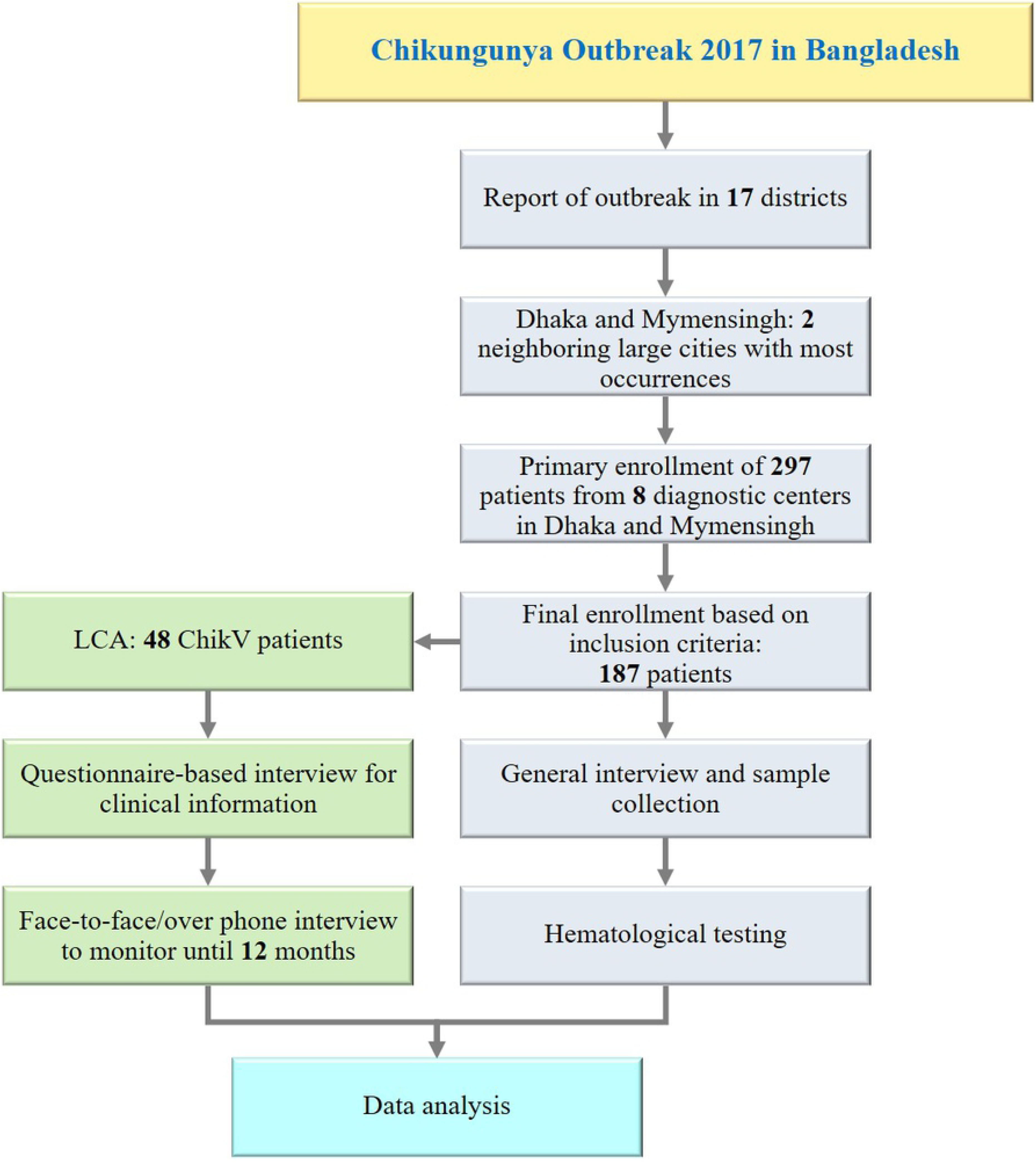
A schematic illustration of key features and work flow for the study.

### Ethics Statement

The methodology and protocol used for this study was reviewed and endorsed by the Graduate Research Ethics Committee, (Headed by the Dean) School of Life Sciences, Shahjalal University of Science and Technology. All participants or their legal representative gave written informed consent according to the Declaration of Helsinki.

### Sero-biochemical and long-term effect study

Biochemical and serological test results were collected after they were prescribed and performed by a specialist physician and a specialist diagnostic center respectively. Relative intensity of joint pain was evaluated using a numerical rating (NR) scale starting from 0 to 10. A rating of 0 indicated that the individual had no joint pain, and a rating of 10 indicates intolerable joint pain. Using these relative scores given by the patients, the pain intensity was categorized as mild (NR 1 – 4), moderate (NR 5 – 7) and severe (8 – 10). The anatomical location(s) of the pain and how long it existed after ChikV infection were also documented. For long-term consequences assessment (LCA) of the after-effect of Chikungunya, consented patients were interviewed with a standard questionnaire 2, 4, 6, 9 and 12 months after the viremic phase (M2, M4, M6, M9 and M12) (**S2 Questionnaire)**. Data were analyzed using SPSS (Statistical Package for Social Sciences) and statistical significance was tested using both one-tailed and two-tailed experiments including Student T-test, z statistic, χ2 test and McNemar tests (for matched pairs of subjects) at a 99% (p = 0.01) and 95% (p = 0.05) confidence level.

## Results

### Features of the patient’s cohort

Among the 187 confirmed (using RT-PCR and/or immunological techniques) Chikungunya (**Table** 1) patients, 117 (62.6%) patients were from the Dhaka region, while 70 (37.4%) were from Mymensingh. Interestingly, 18 patients from Mymensingh reported that they travelled to Dhaka in the weeks before the inclusion. The age range of ChikV positive patients was between 3 and 84 years, with the majority of cases involving the age group 41-59 years (**Fig** 2). Also, 32 children (**≤15 yrs)** were included in this study.

**Fig 2:**
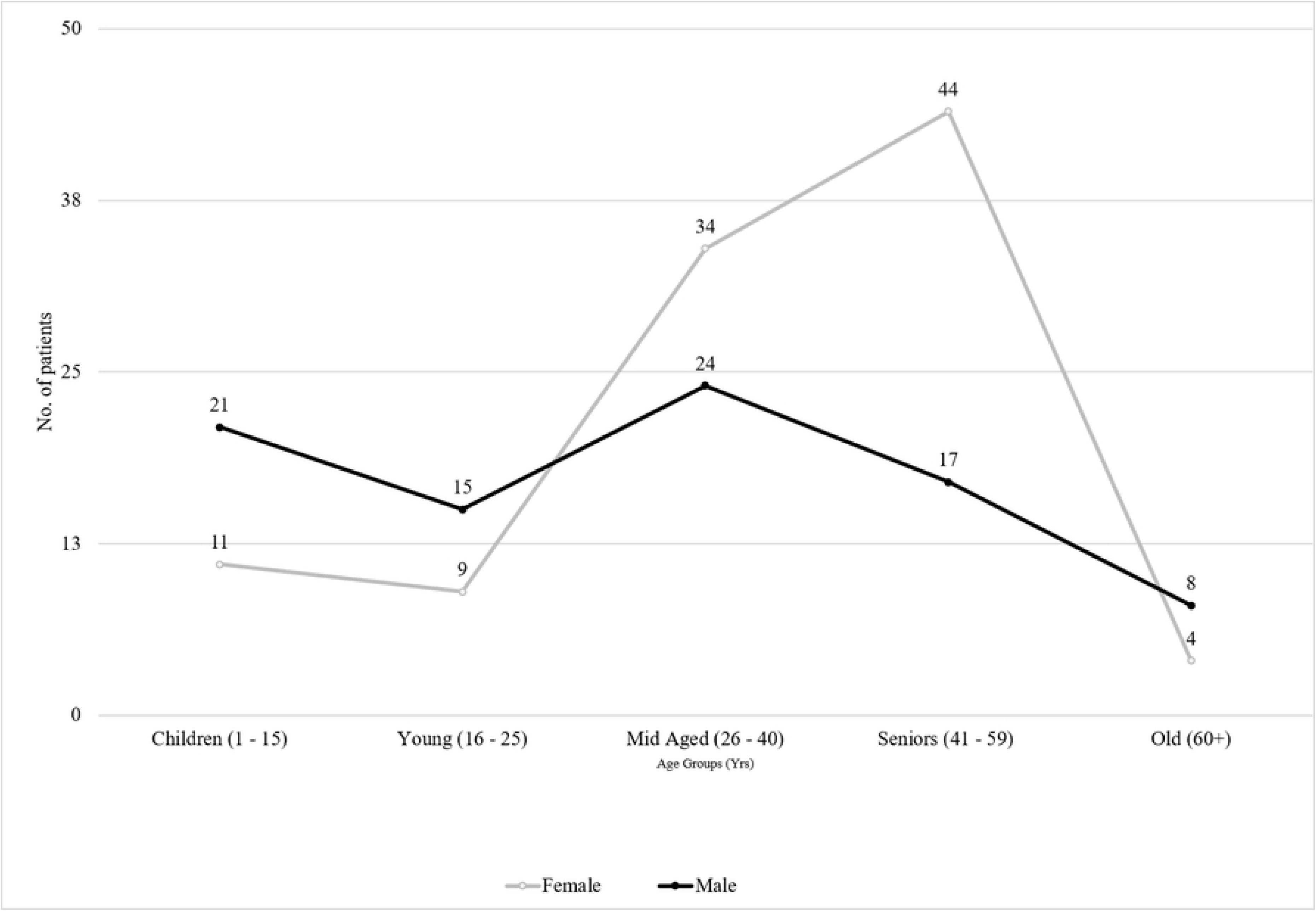
Age and gender distribution of the sample pool.

**Table 1:** Diagnostic outcomes of sero-samples

| Parameter | Finally enrolled patients<br>n (%) | LCA group<br>n (%) |
| --- | --- | --- |
| Total number of patients | 187 (100%) | 47 (100) |
| By Immunochromatography | 135 (72.2%) |  |
| By IgM ELISA | 48 (25.7%) |  |
| By both immunochromatography and IgM ELISA | 4 (2.1%) |  |

*LCA: Long-term consequence assessment; n: number of respondents*

### Demographic Data

Randomly collected samples and demographic data analysis revealed that females were more prone to Chikungunya (M:F=1:1.34) (**Table** 2). Among 187 patients, 2 were admitted to a hospital and most of the patients visited doctors and/or a diagnostic center within, on average, 5.1 days after the first symptoms.

**Table 2:** Demographic features of ChikV positive patients

| Characteristics | Finally enrolled patients<br>Value | LCA group<br>Value |
| --- | --- | --- |
| Male: Female (Ratio) | 1:1.34 | 1:1.04 |
| Mean years of age ± Standard Deviation (Female) | 38.5 ± 15.56 | 33.42 ± 8.52 |
| Mean years of age ± Standard Deviation (Male) | 29 ± 17.58 | 33.44 ± 11.42 |
| Time from onset to diagnostic center visit (Mean days ± Standard Deviation) | 5.1 ± 2.9 | 4.8 ± 2.6 |
| Hospitalization | 2 (0.54%) | 0 (0.0%) |

*LCA: Long-term consequence assessment*

### Signs, Symptoms and Clinical Features

The symptoms of ChikV infected patients are presented in **Table** 3. The most common feature of the ChikV infection was high fever (39.88°C on average) and arthralgia, found to be present in ∼98% of patients. Arthralgic pain was more frequently reported between day 1 to 3 in the infected persons, while fever was more prominent at day 4 or 5, myalgia between days 4 to 6, skin rashes between 6 to 7 days and itching on day 7. Other noticeable symptoms included swelling, stiffness and redness of joints, itching, headache, cough, insomnia, fatigue and dizziness.

**Table 3:**
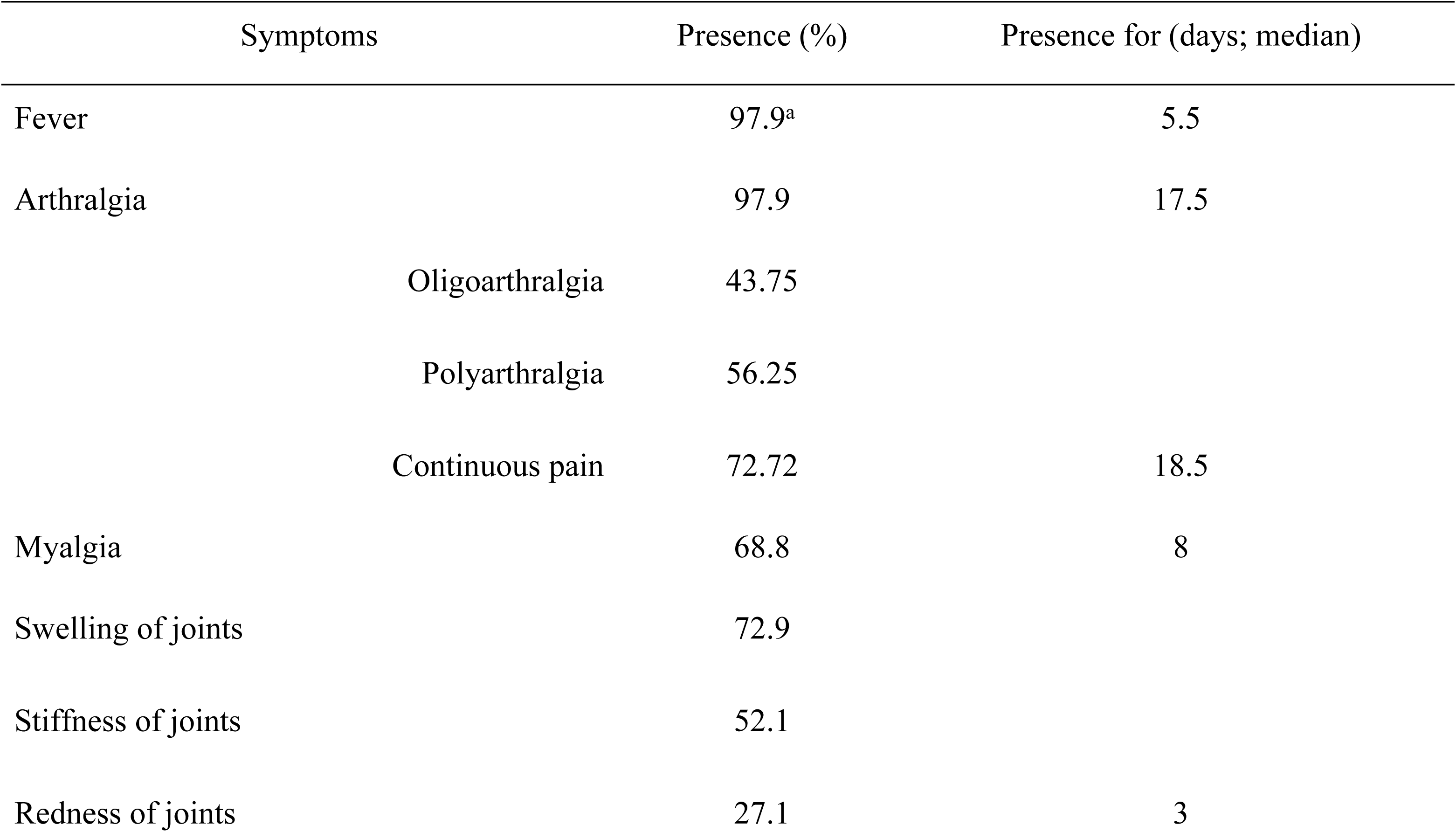
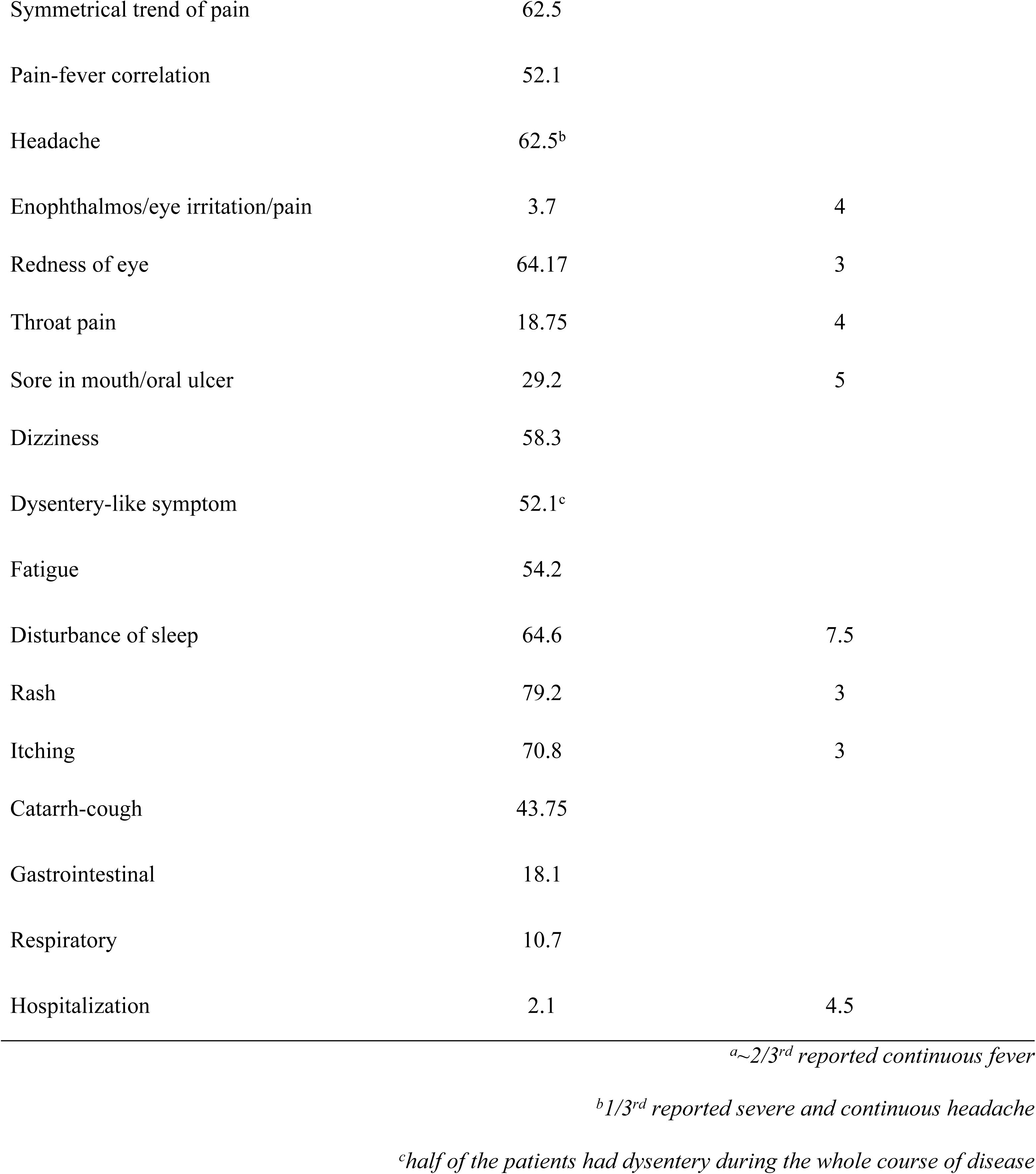
Signs and symptoms recorded from ChikV positive patients during acute phase

| Symptoms | Presence (%) | Presence for (days; median) |
| --- | --- | --- |
| Fever | 97.9<sup>a</sup> | 5.5 |
| Arthralgia | 97.9 | 17.5 |
| Oligoarthralgia | 43.75 |  |
| Polyarthralgia | 56.25 |  |
| Continuous pain | 72.72 | 18.5 |
| Myalgia | 68.8 | 8 |
| Swelling of joints | 72.9 |  |
| Stiffness of joints | 52.1 |  |
| Redness of joints | 27.1 | 3 |
| Symmetrical trend of pain | 62.5 |  |
| Pain-fever correlation | 52.1 |  |
| Headache | 62.5<sup>b</sup> |  |
| Enophthalmos/eye irritation/pain | 3.7 | 4 |
| Redness of eye | 64.17 | 3 |
| Throat pain | 18.75 | 4 |
| Sore in mouth/oral ulcer | 29.2 | 5 |
| Dizziness | 58.3 |  |
| Dysentery-like symptom | 52.1<sup>c</sup> |  |
| Fatigue | 54.2 |  |
| Disturbance of sleep | 64.6 | 7.5 |
| Rash | 79.2 | 3 |
| Itching | 70.8 | 3 |
| Catarrh-cough | 43.75 |  |
| Gastrointestinal | 18.1 |  |
| Respiratory | 10.7 |  |
| Hospitalization | 2.1 | 4.5 |

<sup>a</sup>~2/3<sup>rd</sup> reported continuous fever
<sup>b</sup>1/3<sup>rd</sup> reported severe and continuous headache
<sup>c</sup>half of the patients had dysentery during the whole course of disease

Arthralgia was observed at 12 different anatomical sites (**Fig** 3), with hand joints (fingers and wrist), leg joints (ankle, knee and feet), and shoulder and neck joints being most often affected in the Chikungunya patients. Importantly, arthralgia was typically symmetrical. The intensity of the pain was stronger when patients tried to move. None of the patients reported arthralgia specific to a single anatomical site. Other signs and symptoms which were less frequent included gastrointestinal (GI) and respiratory (RD) complaints.

**Fig 3:**
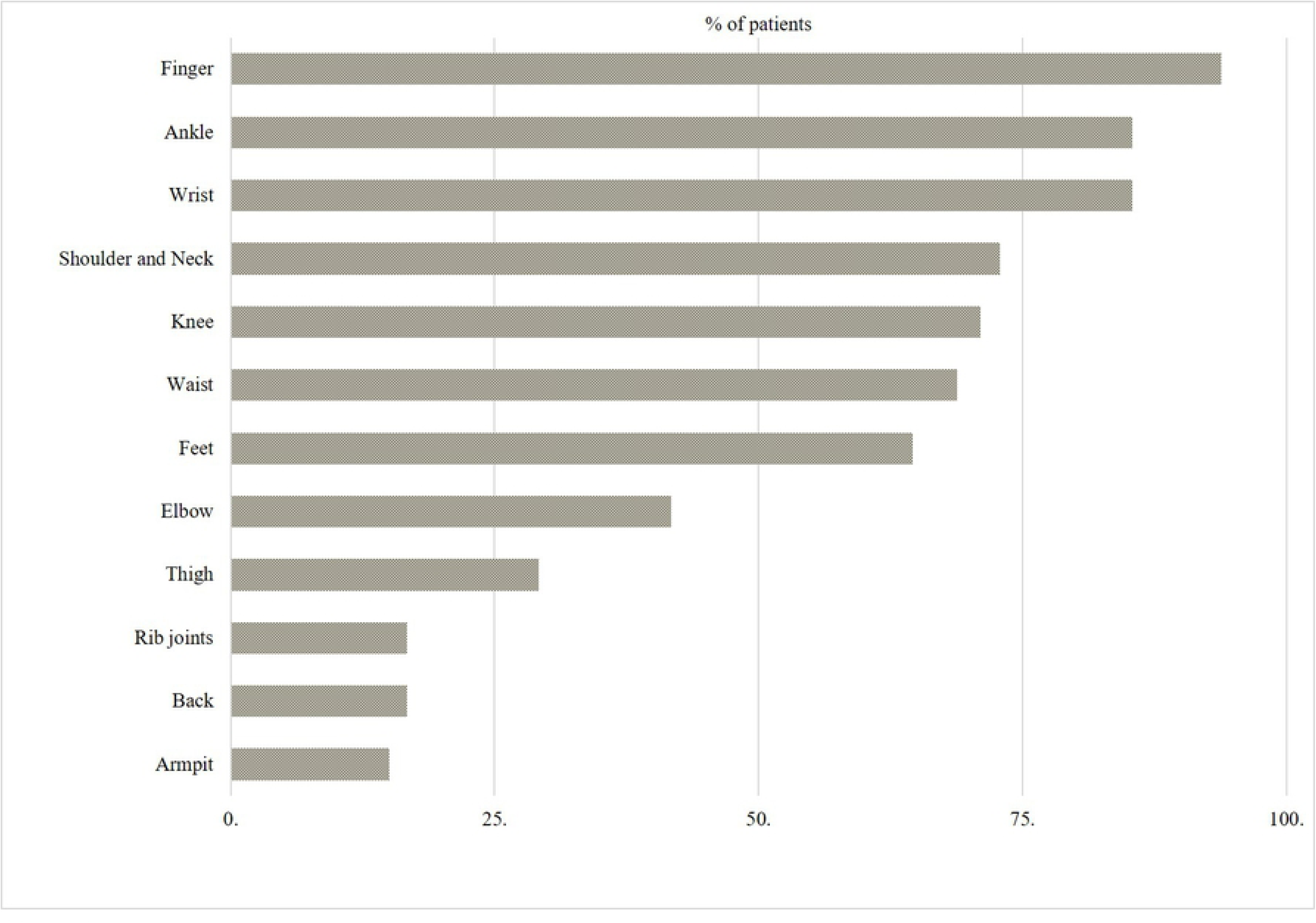
Sites of pain due to Chikungunya infection

The signs and symptoms pattern of Chikungunya seemed remarkably different in children compared to adults (**Table** 4). Arthralgia was less present while vomiting was more frequently reported in children. In addition, the frequency of skin rash was notably higher.

**Table 4:** Differences in clinical manifestations of ChikV positive children and adults

| | Children ($\leq 15$ yrs)<br>(%) | Adults ($> 15$ yrs)<br>(%) | *p* value* |
| --- | --- | --- | --- |
| ***Symptoms at onset (day 0 – day 7)*** |  |  |  |
| Arthralgia | 87.5 | 100 | $< 0.0001$ |
| Headache | 84.4 | 57.4 | $< 0.0001$ |
| Rash | 37.5 | 87.7 | $< 0.0001$ |
| Itching | 43.75 | 76.1 | $< 0.0001$ |
| Vomiting | 64.25 | 25 | $< 0.0001$ |
| ***Symptoms at day 60*** |  |  |  |
| Arthralgia | 15.6 | 37.5 | $< 0.0001$ |
| Headache | 00.0 | 8.33 | $< 0.0001$ |
| Vomiting/vomiting tendency | 00.0 | 10.4** | $< 0.0001$ |

\*Calculated from *t*-test
\*\*All respondents were females

### Hematological Findings

Hematological analysis of ChikV positive patients revealed that hemoglobin level was significantly low in both children and adults compared to standard reference value (**Table** 5). For ChikV positive patients, the complete white blood cell (WBC) counts ranged from 2 to 12.6 K/μL, of which neutrophil (NTP) counts ranged between 32-80% and lymphocyte (LPC) counts ranged between 14 – 56%. Platelet counts ranged from 85 K/μL to 547 K/μL. Although the majority of the ChikV positive patients were within normal ranges for whole WBC, neutrophils, LPC and platelets (PLT), many patients represented varying degrees of lymphopenia when compared to reference values (**Fig** 4).

**Fig 4:**
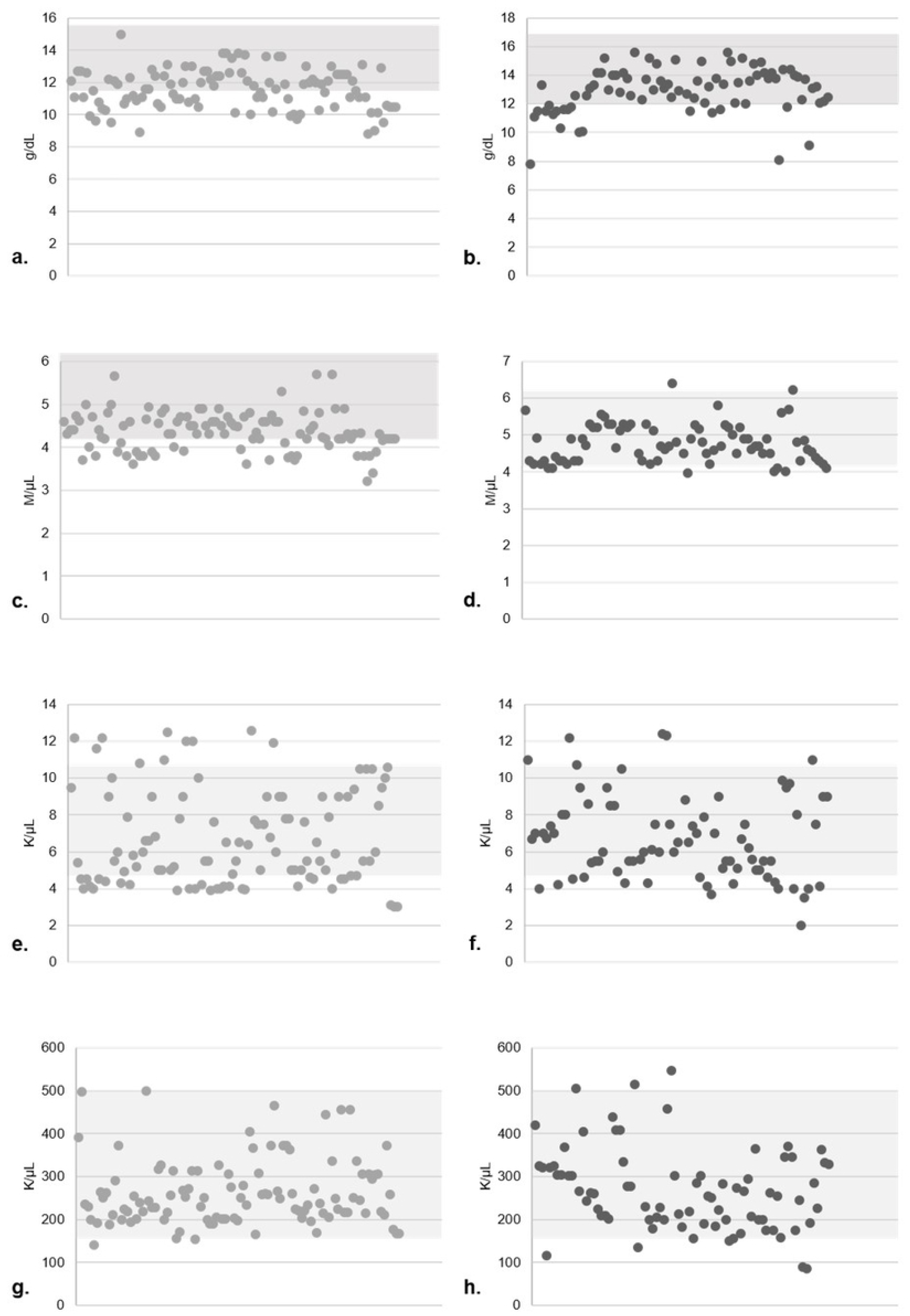
Scatter diagram of hematological findings of ChikV positive patients. Shadowed area denotes the reference ranges. a) Haemoglobin level in females. b) Haemoglobin levels in males. c) RBC counts in females. d) RBC counts in males. e) WBC counts in females. f) WBC counts in males. g) Platelet counts in females. h) Platelet counts in males

**Table 5:**
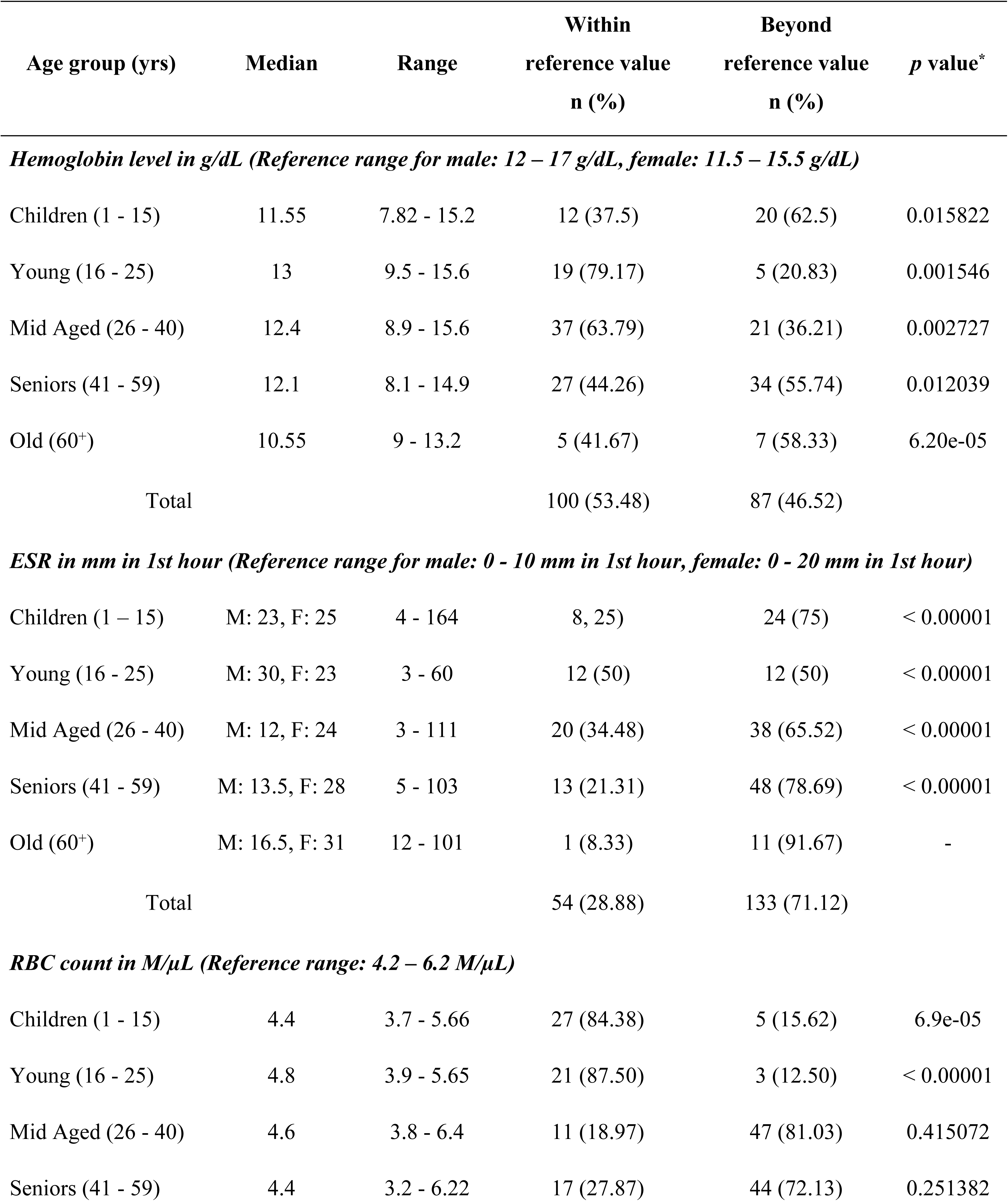
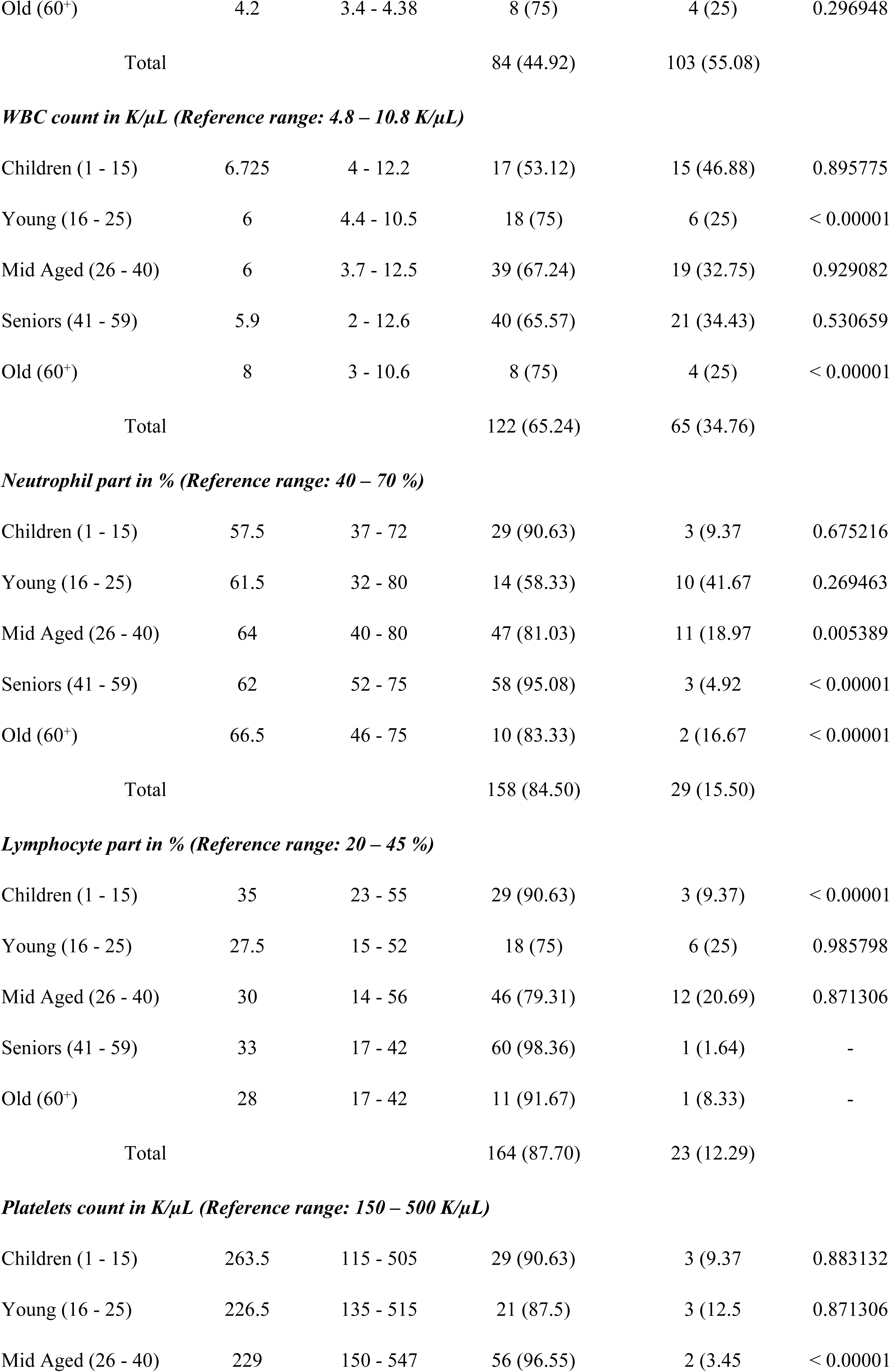
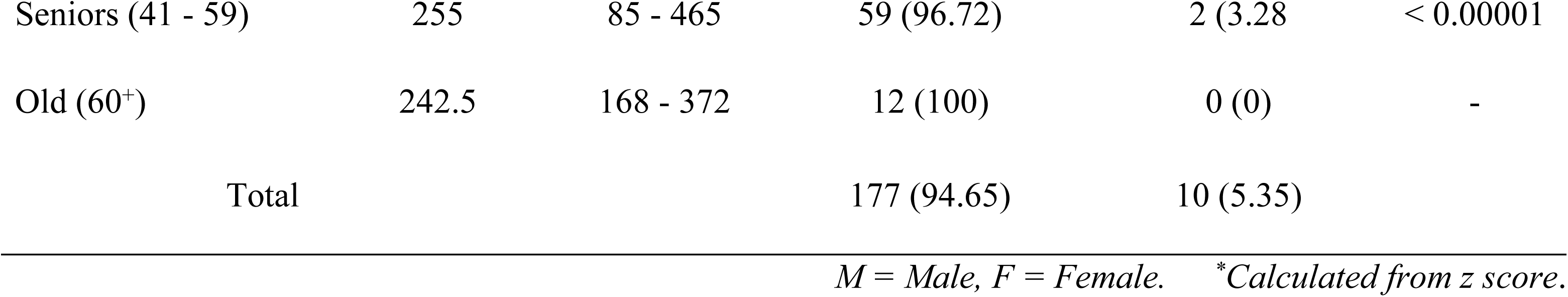
Hematological findings in ChikV positive patients according to age groups

| Age group (yrs) | Median | Range | Within<br>reference value<br>n (%) | Beyond<br>reference value<br>n (%) | p value* |
| --- | --- | --- | --- | --- | --- |
| **Hemoglobin level in g/dL (Reference range for male: 12 – 17 g/dL, female: 11.5 – 15.5 g/dL)** |  |  |  |  |  |
| Children (1 - 15) | 11.55 | 7.82 - 15.2 | 12 (37.5) | 20 (62.5) | 0.015822 |
| Young (16 - 25) | 13 | 9.5 - 15.6 | 19 (79.17) | 5 (20.83) | 0.001546 |
| Mid Aged (26 - 40) | 12.4 | 8.9 - 15.6 | 37 (63.79) | 21 (36.21) | 0.002727 |
| Seniors (41 - 59) | 12.1 | 8.1 - 14.9 | 27 (44.26) | 34 (55.74) | 0.012039 |
| Old (60<sup>+</sup>) | 10.55 | 9 - 13.2 | 5 (41.67) | 7 (58.33) | 6.20e-05 |
| Total |  |  | 100 (53.48) | 87 (46.52) |  |
| **ESR in mm in 1st hour (Reference range for male: 0 - 10 mm in 1st hour, female: 0 - 20 mm in 1st hour)** |  |  |  |  |  |
| Children (1 – 15) | M: 23, F: 25 | 4 - 164 | 8, 25) | 24 (75) | < 0.00001 |
| Young (16 - 25) | M: 30, F: 23 | 3 - 60 | 12 (50) | 12 (50) | < 0.00001 |
| Mid Aged (26 - 40) | M: 12, F: 24 | 3 - 111 | 20 (34.48) | 38 (65.52) | < 0.00001 |
| Seniors (41 - 59) | M: 13.5, F: 28 | 5 - 103 | 13 (21.31) | 48 (78.69) | < 0.00001 |
| Old (60<sup>+</sup>) | M: 16.5, F: 31 | 12 - 101 | 1 (8.33) | 11 (91.67) | - |
| Total |  |  | 54 (28.88) | 133 (71.12) |  |
| **RBC count in M/μL (Reference range: 4.2 – 6.2 M/μL)** |  |  |  |  |  |
| Children (1 - 15) | 4.4 | 3.7 - 5.66 | 27 (84.38) | 5 (15.62) | 6.9e-05 |
| Young (16 - 25) | 4.8 | 3.9 - 5.65 | 21 (87.50) | 3 (12.50) | < 0.00001 |
| Mid Aged (26 - 40) | 4.6 | 3.8 - 6.4 | 11 (18.97) | 47 (81.03) | 0.415072 |
| Seniors (41 - 59) | 4.4 | 3.2 - 6.22 | 17 (27.87) | 44 (72.13) | 0.251382 |
| Old (60<sup>+</sup>) | 4.2 | 3.4 - 4.38 | 8 (75) | 4 (25) | 0.296948 |
| Total |  |  | 84 (44.92) | 103 (55.08) |  |
| ***WBC count in K/ $\mu$ L (Reference range: 4.8 – 10.8 K/ $\mu$ L)*** |  |  |  |  |  |
| Children (1 - 15) | 6.725 | 4 - 12.2 | 17 (53.12) | 15 (46.88) | 0.895775 |
| Young (16 - 25) | 6 | 4.4 - 10.5 | 18 (75) | 6 (25) | < 0.00001 |
| Mid Aged (26 - 40) | 6 | 3.7 - 12.5 | 39 (67.24) | 19 (32.75) | 0.929082 |
| Seniors (41 - 59) | 5.9 | 2 - 12.6 | 40 (65.57) | 21 (34.43) | 0.530659 |
| Old (60<sup>+</sup>) | 8 | 3 - 10.6 | 8 (75) | 4 (25) | < 0.00001 |
| Total |  |  | 122 (65.24) | 65 (34.76) |  |
| ***Neutrophil part in % (Reference range: 40 – 70 %)*** |  |  |  |  |  |
| Children (1 - 15) | 57.5 | 37 - 72 | 29 (90.63) | 3 (9.37) | 0.675216 |
| Young (16 - 25) | 61.5 | 32 - 80 | 14 (58.33) | 10 (41.67) | 0.269463 |
| Mid Aged (26 - 40) | 64 | 40 - 80 | 47 (81.03) | 11 (18.97) | 0.005389 |
| Seniors (41 - 59) | 62 | 52 - 75 | 58 (95.08) | 3 (4.92) | < 0.00001 |
| Old (60<sup>+</sup>) | 66.5 | 46 - 75 | 10 (83.33) | 2 (16.67) | < 0.00001 |
| Total |  |  | 158 (84.50) | 29 (15.50) |  |
| ***Lymphocyte part in % (Reference range: 20 – 45 %)*** |  |  |  |  |  |
| Children (1 - 15) | 35 | 23 - 55 | 29 (90.63) | 3 (9.37) | < 0.00001 |
| Young (16 - 25) | 27.5 | 15 - 52 | 18 (75) | 6 (25) | 0.985798 |
| Mid Aged (26 - 40) | 30 | 14 - 56 | 46 (79.31) | 12 (20.69) | 0.871306 |
| Seniors (41 - 59) | 33 | 17 - 42 | 60 (98.36) | 1 (1.64) | - |
| Old (60<sup>+</sup>) | 28 | 17 - 42 | 11 (91.67) | 1 (8.33) | - |
| Total |  |  | 164 (87.70) | 23 (12.29) |  |
| ***Platelets count in K/ $\mu$ L (Reference range: 150 – 500 K/ $\mu$ L)*** |  |  |  |  |  |
| Children (1 - 15) | 263.5 | 115 - 505 | 29 (90.63) | 3 (9.37) | 0.883132 |
| Young (16 - 25) | 226.5 | 135 - 515 | 21 (87.5) | 3 (12.5) | 0.871306 |
| Mid Aged (26 - 40) | 229 | 150 - 547 | 56 (96.55) | 2 (3.45) | < 0.00001 |
| Seniors (41 - 59) | 255 | 85 - 465 | 59 (96.72) | 2 (3.28) | < 0.00001 |
| Old (60<sup>+</sup>) | 242.5 | 168 - 372 | 12 (100) | 0 (0) | - |
| Total |  |  | 177 (94.65) | 10 (5.35) |  |

*M = Male, F = Female. \*Calculated from z score.*

However, at the onset of the disease, no significant correlation was observed between the level of leukocytopenia and the intensity of arthralgic pain. Red blood cell (RBC) count was remarkably above the reference range in the mid aged and senior groups. The interquartile range of the RBC was between 3.2 to 6.22 M/μL and the median value was 4.5 M/μL. Erythrocyte sedimentation rate (ESR) was significantly higher in all age groups, especially in female patients (**Table** 5). Further, a significant correlation was obtained between the different age groups and RBC counts, neutrophil counts and leucocyte counts of the ChikV positive patients (**S3 Table**).

### Characteristics of long-term arthralgia in ChikV infected patients

#### Long-term arthralgia associated with ChikV infection

All patients enrolled in the LCA group were interviewed using a questionnaire at M2, M4, M6, M9 and M12 post-ChikV infection to monitor persistence of fever, arthralgia and other clinical symptoms. None of our monitored patients reported a relapse of the fever. The percentage of patients suffering from long-term arthralgia decreased significantly (till M6) after the acute phase of the infection and then raised to ∼19% at M9 and M12. Most of our enrollees complained of intermittent arthralgia, with successive recovery and relapse; none of the patients complained of permanent arthralgia at any timepoint after the acute phase and all of the respondents reported that the intensity of the pain was significantly reduced after M2. Of note, all of our enrolled patients in the LCA group suffered from arthralgia between day 0 to 7 and none of them suffered from joint pains prior to the ChikV infection. The McNemar test for matched pairs of subjects revealed that the site of arthralgic pain (**S4 Table**) remained the same at each time point. The percentage of patients suffering from myalgia decreased a lot after the acute phase of the infection and stabilized by M6. None of the patients reported to have myalgia at M9 and M12 (**Fig** 5). When ChikV-induced arthralgia relapsed, it was symmetrical, involving more than 2 different anatomical locations. Most frequently, the finger joints, wrists and ankles were affected; regardless of age and sex, some patients complained of pain at elbow joint and knee.

**Fig 5:**
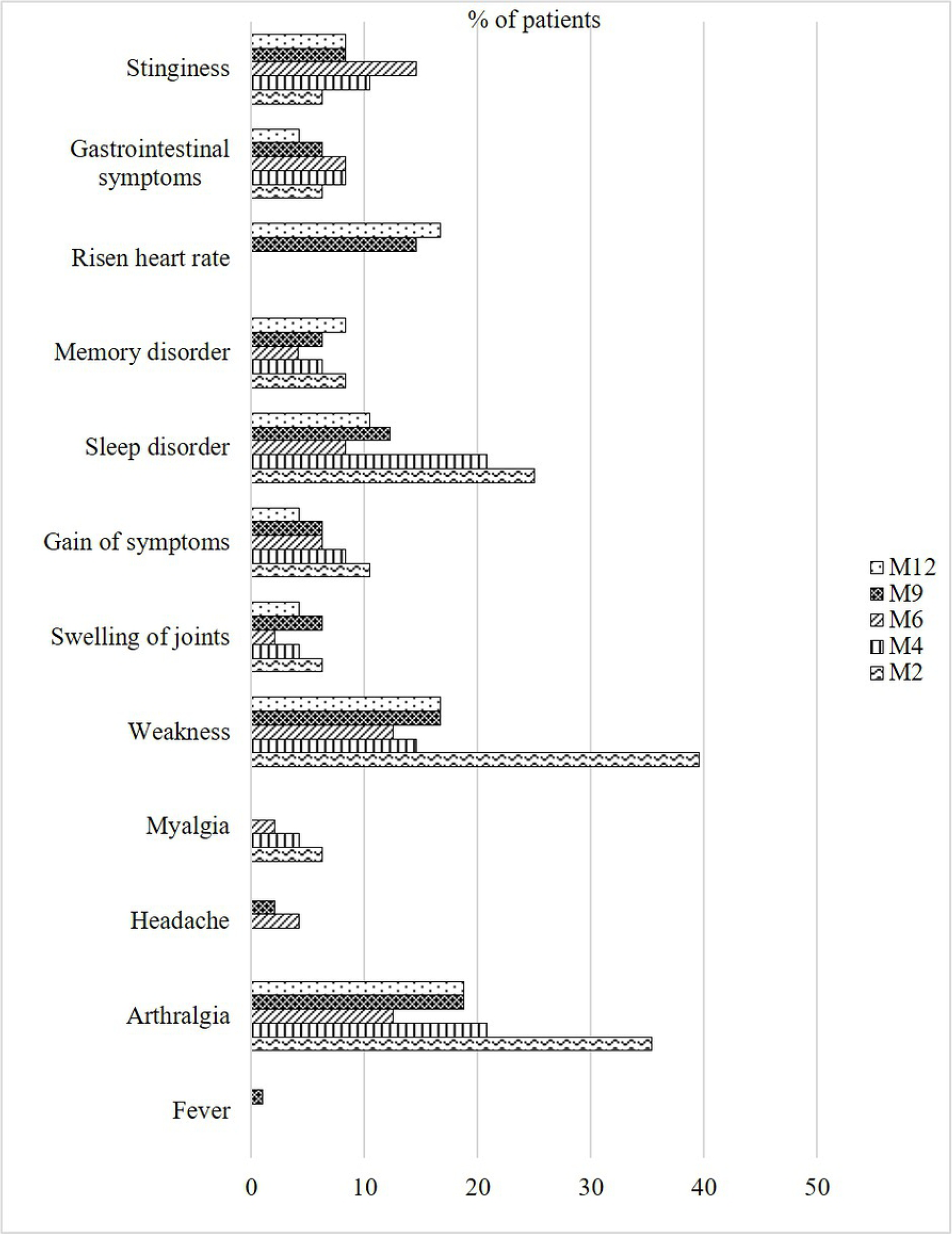
Persistence of Chikungunya symptoms over time course in ChikV positive patients

We noted that the number of sites affected by arthralgia gradually diminished in patients still suffering until M2, with only 23% patients suffering from polyarthralgia. However, the number of anatomical locations further decreased significantly in M4 and M6 and then stabilized at M9 and M12 (50% and 64% respectively, *p* value < 0.0001).

#### Other long-term clinical signs associated with ChikV infection

At M6, M9 and M12, the LCA group displayed other symptoms including local swelling of joints, cutaneous and dermatological symptoms and post-infection weakness. Additionally, sleep, memory and/or concentration disorders as well as depression and stinginess were remarkably associated. Furthermore, between M6 and M12, 16.67% (n = 8) complained that they frequently suffered tachycardia during working, even though none of them had any previous heart complications.

The number of patients who visited a physician increased significantly between M6 (4; 8.33%) and M12 (16; 33%, data not shown) (p < 0.005). However, the most significant after-effect of the infection in our study population appeared to be post infection weakness. Around 40% of the patients reported to have continuous weakness at M2. The percentage decreased significantly at M6, increasing however again to ∼17% at M9 and M12. Besides, more than 20% of patients complained about sleep disorder at M2 and M4, but the percentage diminished to less than 10% at M6. However, complaints of disturbed sleep slightly increased at M9 and M12 (**Fig** 5).

Over 10% of patients complained of new symptoms at M2 which they had not suffered from during the acute phase of the infection (**Fig** 5). Even at M6, 6.25% patients reported to suffer new symptoms and more than 4% patients to experience new symptoms at M9. However, we did not attempt to find any association between the newly gained symptoms and the effect of the RNA virus infection. Although the new symptoms were generally sleeping problems, swelling of joints, arthritis-like symptoms and memory problems were also reported. Although the respondents did not display any neurological dysfunction, the mild memory problems could not be excluded to result from the ChikV infection. Some of the patients reported that they were frequently prone to depression and partially lost control over their temper.

#### Post infection impact on daily life at M6 and M12

Arthralgia coupled with weakness in patients at M6 and M12 were highly incapacitating for daily life activities, professional life and leisure activities (**Table** 6). Most of the patients having chronic arthralgia complained of pain when rising from sitting and lying, walking or picking up a load. Twenty-seven percent and ∼23% patients respectively in M6 and M12 reported that the arthralgia affected their professional activities. Remarkably, ∼31% patients complained that arthralgia had disturbed them in leisure. Moreover, all the patients having memory problems complained that it had significant impact on their day to day life activities.

**Table 6:**
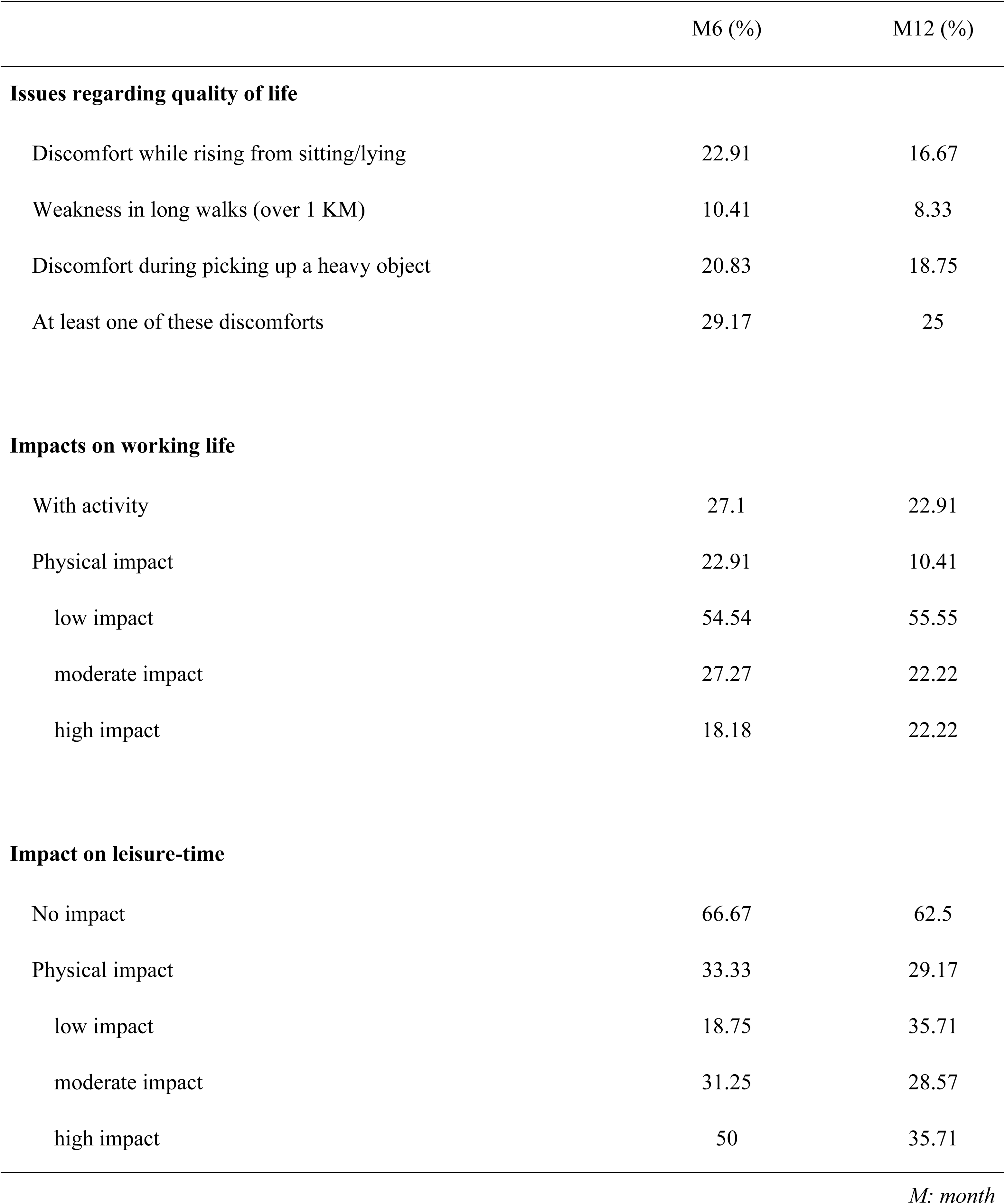
Impact of arthralgia on daily life for patients at M6 and M12

## Discussion

The Chikungunya outbreak of 2017 in Bangladesh appeared as an epidemic manifestation with 23 of 65 districts of the country infected. This study presents clinical and epidemiological data of this Chikungunya outbreak.

Bangladesh is a riverine monsoon country, and as such an ideal vicinity for the emergence of arboviral diseases including Dengue and Chikungunya. As both have overlapping pathophysiological mechanisms and proceed simultaneously, it is real challenge for physicians to distinguish among them, especially during the early stages of infection [30].

ChikV was found to infect all ages and both sexes; however, ratios varied. A higher percentage of cases was observed in adult females (56.7%) than males (38%) and female children (43.3%). This is different from previously reported ratios for the Asian lineage where male cases were more frequent [30-34], except for the reports from Pakistan where females were more prone to be infected by the ChikV [35]. The higher percentage of adult female cases may be due to higher levels of exposure to infected vectors in the home environment, since Bengali women spend more time at home and the mosquitoes are commonly found indoors [36-39]. In addition, mosquitoes can trace estrogen which works as a lure and since females exhibit higher levels of estrogen, mosquitoes tend to bite them more [40-41]. The difference in number of cases in the age groups may not reflect vulnerability of any specific group but indicate the general population structure in the country [42], i.e. the infection trend was not biased to any age group.

Irrespective of sex, the combination of fever and severe arthralgia (present in 97.9% of cases) can be regarded as the cardinal hallmark of the Chikungunya 2017 outbreak in Bangladesh. This is consistent with the previous outbreak report (83.3 to 98%), though the values were less in case of the children [31, 43-45] (**Table** 2, **Table** 3). However, in an Indian outbreak of the virus in Kerala in 2007, arthralgia was found to be the initial symptom in only ∼17% patients [33].

We found a symmetrical presentation of arthralgia in most of the cases (62.5%), while a higher percentage of patients reported polyarthralgia (56.25%) than oligoarthralgia (43.75%). In addition, we observed that finger joints (93.8%) and wrist (85.4%) joints were the most affected sites. In the acute phase, the frequency of incapacitating pain involving certain peripheral joints (**Fig** 3) was found to be comparable with the study of Queyriaux et al. 2008 and Staikowsky et al. 2008; however, it contrasted with earlier reports from India and Suriname [33, 46-49].

Other symptoms including headache, itching, catarrh-cough, dizziness, and dysentery were found to be similar to most of previous studies, except for an unusually high frequency of rash (79.2%), swollen joints (72.9%) and redness of eye (64.17%) in the present study [50].

Over 85% of the patients complained of severe pain with a median NRS score of 9 throughout the acute phase, which was similar as the findings of Staikowsky et al. 2008 [47]. Almost two-thirds (64.6%) of our enrolled patients faced sleep disturbances due to arthralgia and myalgia (**Table** 3). The rate of hospitalization (2.1%) was very low and the outbreak did not cause any fatal outcomes. Nonetheless, the overall severity and the extent of arthralgia-related manifestations suggest that an aggressive strain of Chikungunya virus probably circulated during the outbreak.

The severity of certain clinical manifestations of Chikungunya might depend on several factors including age, gender, immune status, genetic predisposition etc. [51]. Our analysis showed that children (<15 years) had a lower tendency to have skin rash and itching as well as vomiting. Conversely, a significantly higher frequency of headache was observed among the children compared to other age groups. The duration of pain and rate of any relapse of post-infection symptoms were significantly lower among children as compared to other age groups (**Table** 4).

With regards to hematological parameters, distinct markers are yet to be found. Lee et al. (2012) documented several predictable laboratory tests for detecting ChikV, e.g. a drop in lymphocyte count and a higher count of platelets, leukocytes and neutrophils [52]. In our study, a difference in the hematological findings was documented across all age groups. For instance, hemoglobin level was significantly lower in children and older patients; however, RBC counts were significantly above the reference range in mid-aged and senior groups. We were not able to document any significant drop in the lymphocyte counts nor any significant increase in platelets, leucocytes or neutrophils counts. These outcomes are atypical when compared to the reports from the Ahmedabad outbreak, the Caribbean outbreak in Trinidad in 2015, the La Romana outbreak in 2016 and the Kandy outbreak in Sri Lanka in 2006-07 [31, 44, 53-54]. However, the ESR values obtained through our analysis reports a very broad range with significantly elevated rates in most cases across age and sex (p <0.00001) (**Table** 5).

Based on the follow-up of patients with acute ChikV infection who consented to participate, this study shows the evolution of arthralgia, mapping the frequency and location of arthralgic sites during a 12-month time period. Our data reveals that the proportion of patients having ChikV-induced arthralgia decreased at an almost constant rate at each timepoint (**Fig** 5). Myalgia was not a complaint anymore at M6 and thereafter. This is different compared to the higher percentage of patients with long-term symptoms was reported by several studies of Italian and French cohorts of La Re’union Island or metropolitan France [43, 55-57]. Till M12, ChikV-induced arthralgia was mainly symmetrical, and finger joints, wrist, ankle and knee were found to be most affected; this remains consistent with other studies [43, 54].

There is evidence from different countries - especially France, India, Sri Lanka, Malaysia, Colombia, Venezuela and USA - to suggest that sudden rise of heart rate was associated with the infection at both the acute and the chronic phases [58]. In our cohort, cardiovascular manifestations were not reported by patients during the acute phase and till M6. However, 16.67% of the patients experienced abnormal heart rates between M6 and M12. Alvarez et al. underlined the urgent need to explore the cardiovascular impact of a ChikV infection in 2017 [58]. To date, these effects remain to be elucidated.

Weakness during professional activities was noted to be the most prominent after effect of the infection, as almost 40% of our patients reported to have severe weakness at M4. The proportion diminished over time but relapsed several times in some patients till M12. In addition, many patients complained from disturbed sleep, swelling of joints and suffered new symptoms, e.g. memory problems. Although the patients in our study did not display any significant neurological symptoms at the acute phase of disease, we were unable to exclude that these memory problems during the chronic phase resulting from ChikV spread in the central nervous system, as it had been reported that ChikV disseminates to the central nervous system in humans and in animals [59-62]. As was obvious in other studies, chronic ChikV induced complications are considered incapacitating for daily life tasks and impact professional activities and quality of life [56-57].

While the previous studies concerning Chikungunya Outbreak 2017 in Bangladesh were limited within the samples recruited from Dhaka only, this study represents a diverse sample population [30, 51]. In addition, our study was extended to the hematological and chronic outcomes of the outbreak rather than to be confined only within the study of clinical and quality of life parameters [30]. However, this study is not free from any limitations. This recruited only the clinically confirmed cases of Chikungunya, but the studied sample pool was relatively smaller than the previous study [30, 51]. Hossain et al., 2018 reported that, the representation of clinically confirmed cases of ChikV was very low during the 2017 outbreak in Bangladesh due to the high cost of testing and scarcity of diagnostic facilities [30], which might be an explanation behind the recruitment of this smaller sample pool. Moreover, data regarding the clinical, chronic impact and daily life related parameters were collected through retrospective technique, which might be prone to recall bias. Further, it is not unlikely that some respondents have overvalued some clinical symptoms due to the psychological impacts of massive social media coverage of the outbreak. But this study was conducted during the very peak of the outbreak and the patients were monitored and interviewed rigorously at regular intervals, we assume the recall bias was minimized.

In summary, this study alludes to the clinical and epidemiological characteristics of the Chikungunya outbreak of 2017 in Bangladesh. It facilitates our comprehension of the pathophysiology of the disease across age groups and its chronic consequences till M12, a prerequisite for the development of efficient management and therapeutic strategies and for assessing the damage inflicted upon the population by a Chikungunya outbreak.

## Author Contributions

**Conceived and designed the study:** MH. **Questionnaire preparation:** MH, SA. **Survey:** MK, SA, JT. **Follow-up data collection:** SA, JM**. Data curation:** MK, SA, JT. **Formal analysis:** MU, SA, JT. **Contributed reagents/materials/analysis tools:** MH, MU, SA, JT. **Data visualization:** SA. **Script writing:** SA. **Drafted the manuscript:** MH, SA, JT, OV. **Finally approved the manuscript:** All authors.

## Acknowledgments

The authors praise the cooperation by Mr. Mohammad Nazmul Huda of the Uttara Crescent Hospital, Uttara, Dhaka as well as the authorities of the other seven participating diagnostic centers for their helping hands during the patient enrollment. The authors are grateful for the invaluable cooperation of the participants for having confidence and trust in the interviewers and sharing sensitive information with the research team. Special thanks to Ms. Tohura Tahsin for her efforts to improve the quality of our manuscript. The authors would like to thank Dr. Md. Shakhinur Islam Mondal (Associate Professor, Department of Genetic Engineering and Biotechnology, Shahjalal University of Science and Technology) for his kind assistance.

## Supplementary Material

1. S1 Checklist: STROBE Checklist (.docx file)
2. S2 Questionnaire: Questionnaire form used for the LCA (.docx file)
3. S3 Table: (Correlation between age and hematological data
4. S4 Table: A) Sites of pain at different time points. B) Outcomes of McNemar test.

## Data Availability Statement

All supplementary data are given in the Supplementary Material Section. Any additional datasets from this study can be obtained upon request to the corresponding author.

## References

[1] Brighton SW, Prozesky OW, De La Harpe AL. Chikungunya virus infection-A retrospective study of 107 cases. South African Medical Journal. 1983;68(9):313–5.

[2] Pialoux G, Gaüzère BA, Jauréguiberry S, Strobel M. Chikungunya, an epidemic arbovirosis. The Lancet infectious diseases. 2007 May 1;7(5):319–27. doi:10.1016/s1473-3099(07)70107-x.

[3] Thiberville SD, Moyen N, Dupuis-Maguiraga L, Nougairede A, Gould EA, Roques P, de Lamballerie X. Chikungunya fever: epidemiology, clinical syndrome, pathogenesis and therapy. Antiviral research. 2013 Sep 1;99(3):345–70. doi:10.1016/j.antiviral.2013.06.009.

[4] Patterson J, Sammon M, Garg M. Dengue, Zika and chikungunya: emerging arboviruses in the New World. Western Journal of Emergency Medicine. 2016 Nov;17(6):671. doi:10.5811/westjem.2016.9.30904.

[5] Keller, D. M. Debilitating Chikungunya Virus Hits the US. Medscape. 2014. Available: https://www.medscape.com/viewarticle/833612 [Accessed December 2018].

[6] Borgherini G, Poubeau P, Staikowsky F, Lory M, Moullec NL, Becquart JP, Wengling C, Michault A, Paganin F. Outbreak of chikungunya on Reunion Island: early clinical and laboratory features in 157 adult patients. Clinical infectious diseases. 2007 Jun 1;44(11):1401–7. doi:10.1086/517537.

[7] Wahid B, Ali A, Rafique S, Idrees M. Global expansion of chikungunya virus: mapping the 64-year history. International Journal of Infectious Diseases. 2017 May 1;58:69–76. doi:10.1016/j.ijid.2017.03.006.

[8] Lebrun G, Chadda K, Reboux AH, Martinet O, Gaüzère BA. Guillain-Barré syndrome after chikungunya infection. Emerging infectious diseases. 2009 Mar;15(3):495. doi:10.3201/eid1503.071482.

[9] Dupuis-Maguiraga L, Noret M, Brun S, Le Grand R, Gras G, Roques P. Chikungunya disease: infection-associated markers from the acute to the chronic phase of arbovirusinduced arthralgia. PLoS neglected tropical diseases. 2012 Mar 27;6(3):e1446. doi:10.1371/journal.pntd.0001446.

[10] Renault P, Solet JL, Sissoko D, Balleydier E, Larrieu S, Filleul L, Lassalle C, Thiria J, Rachou E, de Valk H, Ilef D. A major epidemic of chikungunya virus infection on Reunion Island, France, 2005–2006. The American journal of tropical medicine and hygiene. 2007 Oct 1;77(4):727–31. doi:10.4269/ajtmh.2007.77.727.

[11] Kosasih H, de Mast Q, Widjaja S, Sudjana P, Antonjaya U, Ma’roef C, Riswari SF, Porter KR, Burgess TH, Alisjahbana B, van der Ven A. Evidence for endemic chikungunya virus infections in Bandung, Indonesia. PLoS neglected tropical diseases. 2013 Oct 24;7(10):e2483. doi:10.1371/journal.pntd.0002483.

[12] Gérardin P, Fianu A, Malvy D, Mussard C, Boussaïd K, Rollot O, Michault A, Gaüzere BA, Bréart G, Favier F. Perceived morbidity and community burden after a Chikungunya outbreak: the TELECHIK survey, a population-based cohort study. BMC medicine. 2011 Dec;9(1):5. doi:10.1186/1741-7015-9-5.

[13] Marimoutou C, Ferraro J, Javelle E, Deparis X, Simon F. Chikungunya infection: selfreported rheumatic morbidity and impaired quality of life persist 6 years later. Clinical Microbiology and Infection. 2015 Jul 1;21(7):688–93. doi:10.1016/j.cmi.2015.02.024.

[14] Thiberville SD, Boisson V, Gaudart J, Simon F, Flahault A, De Lamballerie X. Chikungunya fever: a clinical and virological investigation of outpatients on Reunion Island, South-West Indian Ocean. PLoS neglected tropical diseases. 2013 Jan 17;7(1):e2004. doi:10.1371/journal.pntd.0002004.

[15] Ledrans M, Quatresous I, Renault P, Pierre V. Outbreak of chikungunya in the French Territories, 2006: lessons learned. Weekly releases (1997–2007). 2007 Sep 6;12(36):3262. doi:10.2807/esw.12.36.03262-en.

[16] Mavalankar D, Shastri P, Bandyopadhyay T, Parmar J, Ramani KV. Increased mortality rate associated with chikungunya epidemic, Ahmedabad, India. Emerging infectious diseases. 2008 Mar;14(3):412. doi:10.3201/eid1403.070720.

[17] WHO Media Center. Chikungunya. World Health Organization. 2017. Available: https://www.who.int/news-room/fact-sheets/detail/chikungunya [Accessed December 26, 2018].

[18] Brito CA, Teixeira MG. Increased number of deaths during a chikungunya epidemic in Pernambuco, Brazil. Memorias do Instituto Oswaldo Cruz. 2017 Sep;112(9):650–1. doi:10.1590/0074-02760170124.

[19] icddr, b. First identified outbreak of chikungunya in Bangladesh, 2008. Health Sci Bull. 2009 Mar;7(1):1–6.

[20] Khatun S, Chakraborty A, Rahman M, Banu NN, Rahman MM, Hasan SM, Luby SP, Gurley ES. An outbreak of chikungunya in rural Bangladesh, 2011. PLoS neglected tropical diseases. 2015 Jul 10;9(7):e0003907. doi:10.1371/journal.pntd.0003907.

[21] Kabir I, Dhimal M, Müller R, Banik S, Haque U. The 2017 Dhaka chikungunya outbreak. Lancet Infect Dis. 2017 Nov 1;17(1118):30564–9. doi:10.1016/s1473-3099(17)30564-9.

[22] Uddin KS. Chikungunya-yet another mosquito borne epidemic burden for Bangladesh. KYAMC Journal. 2017 Aug 31;8(1):1–3. doi:10.3329/kyamcj.v8i1.33864.

[23] Rahman, Z. Chikungunya hits nearly every family in Dhaka. The Third Pole. 2017.

[24] Kalam, M. A. New Threat: Chikungunya Outbreak. The Independent. 2017. Available: www.theindependentbd.com/printversion/details/104392 [Accessed December 2018].

[25] Chowdhury, K. R. Chikungunya Outbreak Spreads Beyond Bangladeshi Capital: Health Officials. Benar News 24. 2007. Available: benarnews.org/english/news/bengali/bangladesh-health07212017170830.html [Accessed December 2018].

[26] European Centre for Disease Prevention and Control. Cluster of autochtonous chikungunya cases in France. 2017.

[27] Shishir, M. Chikungunya breaks out in epidemic form. Prothom Alo. 2017. Available: en.prothomalo.com/bangladesh/news/153339/Chikungunya-breaks-out-in-epidemic-form [Accessed December 2018].

[28] Huang JH, Yang CF, Su CL, Chang SF, Cheng CH, Yu SK, Lin CC, Shu PY. Imported chikungunya virus strains, Taiwan, 2006–2009. Emerging infectious diseases. 2009 Nov;15(11):1854. doi:10.3201/eid1511.090398.

[29] Melan A, Aung MS, Khanam F, Paul SK, Riaz BK, Tahmina S, Kabir MI, Hossain MA, Kobayashi N. Molecular characterization of chikungunya virus causing the 2017 26 outbreak in Dhaka, Bangladesh. New microbes and new infections. 2018 Jul 1;24:14–6. doi:10.1016/j.nmni.2018.03.007.

[30] Hossain MS, Hasan MM, Islam MS, Islam S, Mozaffor M, Khan MA, Ahmed N, Akhtar W, Chowdhury S, Arafat SY, Khaleque MA. Chikungunya outbreak (2017) in Bangladesh: Clinical profile, economic impact and quality of life during the acute phase of the disease. PLoS neglected tropical diseases. 2018 Jun 6;12(6):e0006561. doi:10.1371/journal.pntd.0006561.

[31] Langsjoen RM, Rubinstein RJ, Kautz TF, Auguste AJ, Erasmus JH, Kiaty-Figueroa L, Gerhardt R, Lin D, Hari KL, Jain R, Ruiz N. Molecular virologic and clinical characteristics of a chikungunya fever outbreak in La Romana, Dominican Republic, 2014. PLoS neglected tropical diseases. 2016 Dec 28;10(12):e0005189. doi:10.1371/journal.pntd.0005189.

[32] Kumar PS, Arjun MC, Gupta SK, Nongkynrih B. Malaria, dengue and chikungunya in India–An update. Indian Journal of Medical Specialties. 2018 Jan 1;9(1):25–9. doi:10.1016/j.injms.2017.12.001.

[33] Vijayakumar KP, Anish TS, George B, Lawrence T, Muthukkutty SC, Ramachandran R. Clinical profile of chikungunya patients during the epidemic of 2007 in Kerala, India. Journal of global infectious diseases. 2011 Jul;3(3):221. doi:10.4103/0974-777x.83526.

[34] Jain J, Nayak K, Tanwar N, Gaind R, Gupta B, Shastri JS, Bhatnagar RK, Kaja MK, Chandele A, Sunil S. Clinical, serological, and virological analysis of 572 chikungunya patients from 2010 to 2013 in India. Clinical Infectious Diseases. 2017 Jul 1;65(1):133–40. doi:10.1093/cid/cix283.

[35] Badar N, Salman M, Ansari J, Ikram A, Qazi J, Alam MM. Epidemiological trend of chikungunya outbreak in Pakistan: 2016–2018. PLoS neglected tropical diseases. 2019 Apr 18;13(4):e0007118.

[36] Brunkard JM, López JL, Ramirez J, Cifuentes E, Rothenberg SJ, Hunsperger EA, Moore CG, Brussolo RM, Villarreal NA, Haddad BM. Dengue fever seroprevalence and risk factors, Texas–Mexico border, 2004. Emerging infectious diseases. 2007 Oct;13(10):1477. doi:10.3201/eid1310.061586.

[37] Raude J, Setbon M. The role of environmental and individual factors in the social epidemiology of chikungunya disease on Mayotte Island. Health & place. 2009 Sep 1;15(3):689–99. doi:10.1016/j.healthplace.2008.10.009.

[38] Fritel X, Rollot O, Gérardin P, Gaüzère BA, Bideault J, Lagarde L, Dhuime B, Orvain E, Cuillier F, Ramful D, Sampériz S. Chikungunya virus infection during pregnancy, Reunion, France, 2006. Emerging infectious diseases. 2010 Mar;16(3):418. doi:10.3201/eid1603.091403.

[39] Salje H, Lessler J, Paul KK, Azman AS, Rahman MW, Rahman M, Cummings D, Gurley ES, Cauchemez S. How social structures, space, and behaviors shape the spread of infectious diseases using chikungunya as a case study. Proceedings of the National Academy of Sciences. 2016 Nov 22;113(47):13420–5. doi:10.1073/pnas.1611391113.

[40] Lindsay S, Ansell J, Selman C, Cox V, Hamilton K, Walraven G. Effect of pregnancy on exposure to malaria mosquitoes. The Lancet. 2000 Jun 3;355(9219):1972. doi:10.1016/s0140-6736(00)02334-5.

[41] Fradin MS. Mosquitoes and mosquito repellents: a clinician’s guide. Annals of internal medicine. 1998 Jun 1;128(11):931–40. doi:10.7326/0003-4819-128-11-199806010-00013.28

[42] Bangladesh Bureau of Statistics (BBS). Population distribution and internal migration in Bangladesh. 2007.

[43] Larrieu S, Pouderoux N, Pistone T, Filleul L, Receveur MC, Sissoko D, Ezzedine K, Malvy D. Factors associated with persistence of arthralgia among Chikungunya virusinfected travellers: report of 42 French cases. Journal of Clinical Virology. 2010 Jan 1;47(1):85–8. doi:10.1016/j.jcv.2009.11.014.

[44] Sahadeo N, Mohammed H, Allicock OM, Auguste AJ, Widen SG, Badal K, Pulchan K, Foster JE, Weaver SC, Carrington CV. Molecular characterisation of chikungunya virus infections in Trinidad and comparison of clinical and laboratory features with dengue and other acute febrile cases. PLoS neglected tropical diseases. 2015 Nov 18;9(11):e0004199. doi:10.1371/journal.pntd.0004305.

[45] Mattar S, Miranda J, Pinzon H, Tique V, Bolaños A, Aponte J, Arrieta G, Gonzalez M, Barrios K, Contreras H, Alvarez J. Outbreak of Chikungunya virus in the north Caribbean area of Colombia: clinical presentation and phylogenetic analysis. The Journal of Infection in Developing Countries. 2015 Oct 29;9(10):1126–32. doi:10.3855/jidc.6670.

[46] Queyriaux B, Simon F, Grandadam M, Michel R, Tolou H, Boutin JP. Clinical burden of chikungunya virus infection. The Lancet infectious diseases. 2008 Jan 1;8(1):2–3.

[47] Staikowsky F, Le Roux K, Schuffenecker I, Laurent P, Grivard P, Develay A, MichaultRetrospective survey of Chikungunya disease in Reunion Island hospital staff. Epidemiology & Infection. 2008 Feb;136(2):196–206. doi:10.1017/s0950268807008424.

[48] Chopra A, Anuradha V, Ghorpade R, Saluja M. Acute Chikungunya and persistent musculoskeletal pain following the 2006 Indian epidemic: a 2-year prospective rural community study. Epidemiology & Infection. 2012 May;140(5):842-50.29

[49] van Genderen FT, Krishnadath I, Sno R, Grunberg MG, Zijlmans W, Adhin MR. First chikungunya outbreak in Suriname; clinical and epidemiological features. PLoS neglected tropical diseases. 2016 Apr 15;10(4):e0004625.

[50] Teng TS, Kam YW, Lee B, Hapuarachchi HC, Wimal A, Ng LC, Ng LF. A systematic meta-analysis of immune signatures in patients with acute chikungunya virus infection. The Journal of infectious diseases. 2015 Jan 29;211(12):1925–35. doi:10.1093/infdis/jiv049.

[51] Deeba IM, Hasan MM, Al Mosabbir A, Banna Siam MH, Islam MS, Raheem E, Hossain MS. Manifestations of Atypical Symptoms of Chikungunya during the Dhaka Outbreak (2017) in Bangladesh. The American journal of tropical medicine and hygiene. 2019 Apr 29:tpmd190122.

[52] Lee VJ, Chow A, Zheng X, Carrasco LR, Cook AR, Lye DC, Ng LC, Leo YS. Simple clinical and laboratory predictors of Chikungunya versus dengue infections in adults. PLoS neglected tropical diseases. 2012 Sep 27;6(9):e1786. doi:10.1016/j.ijid.2012.05.885.

[53] Kularatne SA, Gihan MC, Weerasinghe SC, Gunasena S. Concurrent outbreaks of Chikungunya and Dengue fever in Kandy, Sri Lanka, 2006–07: a comparative analysis of clinical and laboratory features. Postgraduate medical journal. 2009 Jul 1;85(1005):342–6. doi:10.1136/pgmj.2007.066746.

[54] Shah PS, Shah ND, Patel AS, Kurtadikar SM, Patel KR, Murarka SM, Shah BS, Rao MV. Outbreak of Chikungunya in Ahmedabad: A Report. Biotechnological Research. 2017 Feb 13;3(2):35-8. Available: biotechnologicalresearch.com/index.php/BR/article/download/56/54

[55] Borgherini G, Poubeau P, Jossaume A, Gouix A, Cotte L, Michault A, Arvin-Berod C, Paganin F. Persistent arthralgia associated with chikungunya virus: a study of 88 adult30 patients on reunion island. Clinical Infectious Diseases. 2008 Aug 15;47(4):469–75. doi:10.1086/590003.

[56] Moro ML, Grilli E, Corvetta A, Silvi G, Angelini R, Mascella F, Miserocchi F, Sambo P, Finarelli AC, Sambri V, Gagliotti C. Long-term chikungunya infection clinical manifestations after an outbreak in Italy: a prognostic cohort study. Journal of Infection. 2012 Aug 1;65(2):165–72. doi:10.1016/j.jinf.2012.04.005.

[57] Couturier E, Guillemin F, Mura M, Léon L, Virion JM, Letort MJ, De Valk H, Simon F, Vaillant V. Impaired quality of life after chikungunya virus infection: a 2-year followup study. Rheumatology. 2012 Mar 16;51(7):1315–22. doi:10.1093/rheumatology/kes015.

[58] Alvarez MF, Bolívar-Mejía A, Rodriguez-Morales AJ, Ramirez-Vallejo E. Cardiovascular involvement and manifestations of systemic Chikungunya virus infection: a systematic review. F1000Research. 2017;6. doi:10.12688/f1000research.11078.2.

[59] Barr KL, Khan E, Farooqi JQ, Imtiaz K, Prakoso D, Malik F, Lednicky JA, Long MT. Evidence of chikungunya virus disease in Pakistan since 2015 with patients demonstrating involvement of the central nervous system. Frontiers in public health. 2018; 6:186. doi:10.3389/fpubh.2018.00186.

[60] Couderc T, Chrétien F, Schilte C, Disson O, Brigitte M, Guivel-Benhassine F, Touret Y, Barau G, Cayet N, Schuffenecker I, Desprès P. A mouse model for Chikungunya: young age and inefficient type-I interferon signaling are risk factors for severe disease. PLoS pathogens. 2008 Feb 15;4(2):e29. doi:10.1371/journal.ppat.0040029.

[61] Labadie K, Larcher T, Joubert C, Mannioui A, Delache B, Brochard P, Guigand L, Dubreil L, Lebon P, Verrier B, de Lamballerie X. Chikungunya disease in nonhuman primates involves long-term viral persistence in macrophages. The Journal of clinical investigation. 2010 Mar 1;120(3):894–906. doi:10.1172/jci40104.

[62] Economopoulou A, Dominguez M, Helynck B, Sissoko D, Wichmann O, Quenel P, Germonneau P, Quatresous I. Atypical Chikungunya virus infections: clinical manifestations, mortality and risk factors for severe disease during the 2005–2006 outbreak on Reunion. Epidemiology & Infection. 2009 Apr;137(4):534–41. doi:10.1017/s0950268808001167.

